# Isolation of a potently neutralizing and protective human monoclonal antibody targeting yellow fever virus

**DOI:** 10.1101/2022.02.28.482437

**Authors:** Michael P. Doyle, Joseph R. Genualdi, Adam L. Bailey, Nurgun Kose, Christopher Gainza, Jessica Rodriguez, Kristen M. Reeder, Christopher A. Nelson, Prashant N. Jethva, Rachel E. Sutton, Robin G. Bombardi, Michael L. Gross, Justin G. Julander, Daved H. Fremont, Michael S. Diamond, James E. Crowe

**Affiliations:** Department of Pathology, Microbiology and Immunology, Vanderbilt University Medical Center, Nashville, Tennessee, 37232, USA; Department of Pathology & Immunology, Washington University School of Medicine, St. Louis, MO 63110, USA; The Vanderbilt Vaccine Center, Vanderbilt University Medical Center, Nashville, Tennessee, 37232, USA; Department of Chemistry, Washington University in St. Louis, Saint Louis, MO 63130, USA; Institute for Antiviral Research, Department of Animal, Dairy, and Veterinary Sciences, Utah State University, Logan, Utah, 84322, USA; Department of Biochemistry & Molecular Biophysics, Washington University School of Medicine, St. Louis, MO 63110, USA; Department of Molecular Microbiology, Washington University School of Medicine, St. Louis, MO 63110, USA; Department of Medicine, Washington University School of Medicine, St. Louis, MO 63110, USA; Department of Pediatrics, Vanderbilt University Medical Center, Nashville, Tennessee, 37232, USA

## Abstract

Yellow fever virus (YFV) causes sporadic outbreaks of infection in South America and sub-Saharan Africa. While live-attenuated yellow fever virus vaccines based on three substrains of 17D are considered some of the most effective vaccines in use, problems with production and distribution have created large populations of unvaccinated, vulnerable individuals in endemic areas. To date, specific antiviral therapeutics have not been licensed for human use against YFV or any other related flavivirus. Recent advances in monoclonal antibody (mAb) technology have allowed for identification of numerous candidate therapeutics targeting highly pathogenic viruses, including many flaviviruses. Here, we sought to identify a highly neutralizing antibody targeting YFV envelope (E) protein as a therapeutic candidate. We used human B cell hybridoma technology to isolate mAbs from the circulating memory B cells from human YFV vaccine recipients. These antibodies bound to recombinant YFV E protein and recognized at least five major antigenic sites on E. Two mAbs (designated YFV-136 and YFV-121) recognized a shared antigenic site and neutralized the YFV 17D vaccine strain *in vitro*. YFV-136 also potently inhibited infection by multiple wild-type YFV strains, in part, at a post-attachment step in the virus replication cycle. YFV-136 showed therapeutic protection in two animal models of YFV challenge including hamsters and immunocompromised mice engrafted with human hepatocytes. These studies define features of the antigenic landscape on YFV E protein recognized by the human B cell response and identify a therapeutic antibody candidate that inhibits infection and disease caused by highly virulent strains of YFV.

## Introduction

Yellow fever virus (YFV), the prototype and namesake member of the family *Flaviviridae*, is a historically important human pathogen. Yellow fever disease has been described in the New World since the 1600s, and YFV was first identified in 1927 [1]. The virus has caused numerous epidemics of human disease throughout the world. According to the World Health Organization, 47 countries in Africa and Central and South America currently have regions endemic for yellow fever, and the burden of yellow fever disease during 2019 was as high as 109,000 severe cases and 51,000 deaths [2]. Approximately thirty percent of infected individuals develop severe disease that includes hemorrhagic complications and multiorgan failure, half of which succumb to the infection [3]. Non-human primates serve as the primary reservoir for YFV, with mosquitoes in the *Haemogogus, Sabethes*, and *Aedes* genera serving as the vectors responsible for reservoir maintenance and spillover into humans, typically when humans encroach on primates’ natural ecosystems, in what is referred to as the “sylvatic cycle.” Once within the human population, YFV is spread by a different vector, the anthropophilic *Aedes aegyptii* mosquito, in an “urban cycle” [4]. Beginning in 2018, YFV epidemics began approaching coastal urban centers like Sao Paolo and Rio de Janeiro, sparking concerns that more severe YFV epidemics may occur in the future [5].

YFV is an enveloped virus with a positive-sense, single-stranded RNA genome. The YFV genome is translated as a single polyprotein, which is post-translationally cleaved by a combination of host and viral proteins into 3 structural (pr/M, E, C) and 7 non-structural (NS1, NS2A, NS2B, NS3, NS4A, NS4B, NS5) proteins [6]. The envelope (E) protein is the primary surface-exposed protein on mature particles and is the principal target of the protective humoral immune system [7]. The E protein is comprised of three domains (domain I (DI), DII, and DIII). DII contains several immunodominant epitopes, including the fusion loop (FL), which is a hydrophobic peptide that mediates fusion of viral and host membranes in the late endosome. Domain III on E contains the putative cellular attachment domain [8]. While several attachment factors have been postulated, specific entry receptors for YFV have not yet been identified. The virus enters host cells by receptor-mediated endocytosis, as the low pH of late endosomes triggers conformational changes in the E protein. These changes expose the FL, which inserts into the endosomal membrane, allowing penetration of the RNA genome into the host cytoplasm for translation and replication. Although E protein is the primary target of neutralizing antibodies [9–12], nonstructural 1 (NS1) proteins also can elicit protective antibodies [13–15]. Conversely, prM antibodies may confer the risk of antibody-dependent enhancement of infection by otherwise poorly infectious immature virions [16–18].

The YFV vaccine, based on a strain known as 17D, was created by serial passage and attenuation in the 1930s by Max Theiler [19] and is considered one of the most successful vaccines ever created. However, production of 17D has changed little since its inception, resulting in a system of manufacturing and distribution that have been unable to keep pace with demand: ~400 million people in endemic areas still require vaccination to achieve the herd-immunity threshold required to prevent urban spread of YFV [20]. Fractional dosing has been explored in outbreak settings when vaccine supply is insufficient, but its consequences on generation of durable, long-lasting protection are unknown [21, 22]. YFV vaccine shortages stem principally from the limitations inherent in the legacy methods of vaccine strain propagation still being used. When outbreaks do occur in the setting of vaccine insufficiency, specific licensed antiviral treatments targeting YFV are not available.

Recently, potent neutralizing mAbs against many viral targets have shown efficacy as potential treatments of highly pathogenic agents, including other flaviviruses. Several such antibodies targeting YFV have been described. A mAb designated A5 was identified using phage display technology and showed efficacy in an immunodeficient YFV-17D challenge model [23]. A humanized mAb designated 2C9 showed benefit in hamsters against Jimenez strain [24] and in AG129 mice against YFV 17D-204 challenge [25], supporting the proof of principle for antibodies as a medical countermeasure for YFV. Fully human mAbs with native heavy and light chain pairing, however, are preferred for use in human therapy. A human antibody designated TY014 has been tested in a Phase 1 trial [26], and recently other groups have reported the isolation of human anti-YFV mAbs [27, 28]. Here, we isolated a panel of fully human mAbs targeting E protein to identify candidate therapeutic antibodies. Competition-binding studies mapped these antibodies to several antigenic sites, one of which elicits antibodies that neutralize YFV. *In vitro* studies of the most potent neutralizing mAb, designated YFV-136, revealed that this antibody exerts its neutralizing activity at least partially at a post-attachment step via binding to DII on YFV E protein. Hydrogen-deuterium exchange mass spectrometry (HDX-MS) and neutralization escape virus selection established a key binding and functional epitope for YFV-136 in DII of the E protein. Passive transfer of YFV-136 mAb protected against lethal YFV challenge in a therapeutic setting in two small animal models, Syrian golden hamsters and immunocompromised mice engrafted with human hepatocytes. These studies identify a potently neutralizing mAb targeting YFV and pave the way for further development of this human mAb YFV-136 as a possible candidate therapeutic agent.

## Results

### Isolation of mAbs from YFV vaccine recipients

Peripheral blood mononuclear cells (PBMCs) from four subjects who received a YFV vaccine previously (varying from months to years prior) were transformed *in vitro* with Epstein-Barr virus (EBV) to screen for YFV-reactive antibodies secreted by transformed memory B cells. We screened cell supernatants for binding to recombinant YFV E protein by ELISA and/or binding to YFV-17D-infected cells by flow cytometry. Cells secreting YFV-reactive antibodies were fused to a myeloma partner to generate hybridoma lines, which were cloned by flow cytometric cell sorting. Antibody was purified from serum-free hybridoma cell line supernatants by affinity chromatography. Using these methods, we isolated 15 mAbs from four YFV-immune subjects. These antibodies bound to recombinant E protein according to ELISA with varying half maximal effective concentrations (EC_50_) for binding ranging from 29 to 15,600 ng/mL (**Figure 1A**). Each of the antibodies isolated was tested for ability to neutralize YFV-17D in a focus reduction neutralization test (FRNT) in Vero cells. While most antibodies did not neutralize YFV-17D infection, two mAbs showed inhibitory activity: YFV-121 was moderately neutralizing, with a half maximal inhibitory concentration of 202 ng/mL and YFV-136 showed exceptional potency, with an IC_50_ below 10 ng/mL (**Figure 1B**).

**Figure 1.**
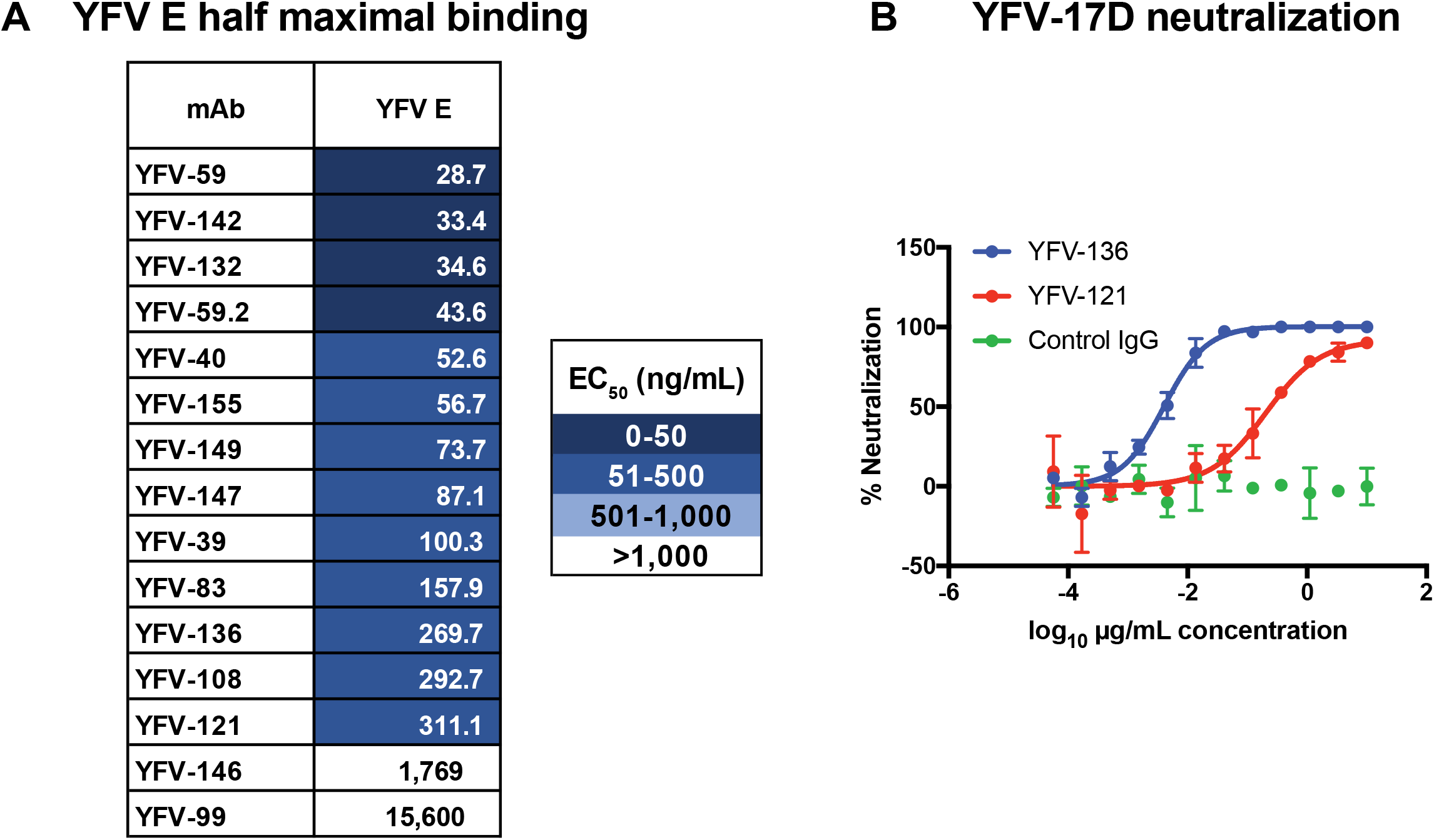
ELISA binding and FRNT neutralization by human mAbs targeting YFV E protein. **A**. Half maximal effective concentrations (EC_50_) of antibody binding to YFV E as determined by ELISA. Values were interpolated using a non-linear regression model in Prism software. Data from a single experiment are shown, representative of three independent experiments. **B**. Focus reduction neutralization test (FRNT) to assess neutralization of YFV-17D by YFV-121 and YFV-136. Neutralization values were fit to a non-linear regression model. Data from a single experiment are shown, representing at least two independent experiments.

### Competition-binding reveals antibodies target several antigenic sites on the E protein

We used biolayer interferometry (BLI) to perform competition-binding studies that enable grouping of antibodies based on the major antigenic sites recognized (**Figure 2A**). In this platform, antigen is loaded onto a biosensor tip, with two antibodies sequentially flowed over the tip. If mAbs recognize non-overlapping antigenic sites, both bind to the coated sensors when applied in sequence. If binding of the first antibody applied to the antigen-coated sensor reduces or prevents binding of the second antibody, the pair of mAbs likely bind to the same or an overlapping antigenic site. We included the previously described pan-flavivirus reactive murine mAb 4G2 targeting the fusion loop (FL) for comparison. The human antibodies recognized six antigenic sites. One group of mAbs, including YFV-39, −40, and −146, competed for binding with 4G2, indicating that these mAbs target regions near the FL epitope on YFV E. The neutralizing mAbs YFV-121 and −136 grouped together, indicating these mAbs target an overlapping antigenic site of neutralization vulnerability on YFV E. YFV-65 also competed for binding to E with YFV-121 and YFV-136, even though it did not neutralize YFV-17D when tested at concentrations as high as 10 μg/mL. These data suggest there are multiple antigenic sites on YFV E, with at least one site being a target of potently neutralizing antibodies.

**Figure 2.**
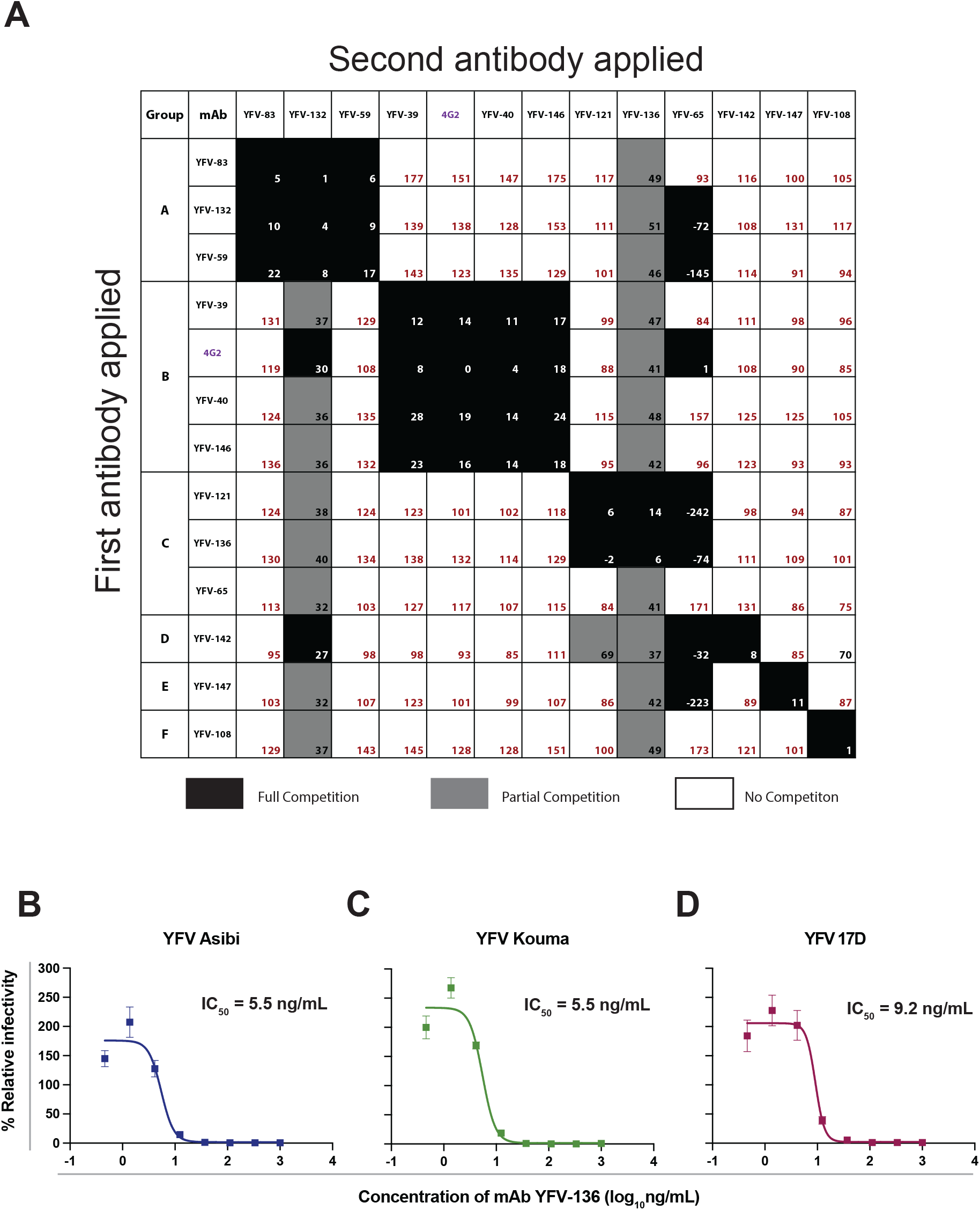
Competition binding and neutralization by human mAbs targeting YFV E protein. **A**. Octet biolayer interferometry competition binding of human mAbs against YFV E. Antibodies listed top to bottom were associated to immobilized YFV E protein, with antibodies shown left to right tested for their ability to bind in the presence of the first antibody. Binding is expressed as a percentage of residual binding, with black boxed indicating compete competition, gray boxed partial competition, and white boxes no competition. Antibodies were clustered based on their competition profiles and labeled A-F. **B-D**. Neutralization of diverse YFV strains assessed by a focus reduction neutralization test (FRNT). Values were fit to a non-linear regression model using Prism software. Three independent experiments were performed in technical triplicate, with data from a single representative experiment shown.

### Neutralization of wild-type YFV strains by mAbs targeting YFV E protein

We next tested YFV-136 for its ability to neutralize wild-type YFV strains under BSL-3 conditions. YFV-136neutralized Asibi and Kouma YFV strains, as well as a different 17D vaccine strain with high potency (**Figure 2B-D**). At lower antibody concentrations, we observed modest enhancement of infectivity in this assay, possibly due to aggregation of virions, as has been seen with other anti-flavivirus antibodies in cells lacking Fcγ receptors [29].

### Identification of antigenic site for mAb YFV-136 using HDX-MS studies

Using HDX-MS, a technique in which antibody binding reduces deuterium labeling of surface-exposed viral protein residues, we identified peptides on the YFV-E protein that that are occluded by binding of YFV-136 Fab fragments (**Figure 3A)**. The start and end residues and the amino acid sequences of the representative E protein peptides that showed differential deuteration in the absence of presence of YFV-136 are shown in **Figure 3B**. The results are summarized in a Woods plot in which the peptides showing a significant decrease in HDX are marked in green lines, and the unaffected peptides are marked as gray solid lines (**Figure 3C**). The HDX protection profile is mapped onto a cartoon representation of the YFV-E dimer (**Figure 3D**). The E protein domain II (DII) near the fusion loop showed the most protection following YFV-136 Fab binding. Some protection against deuteration also was observed in the dimer interface and in DIII.

**Figure 3.**
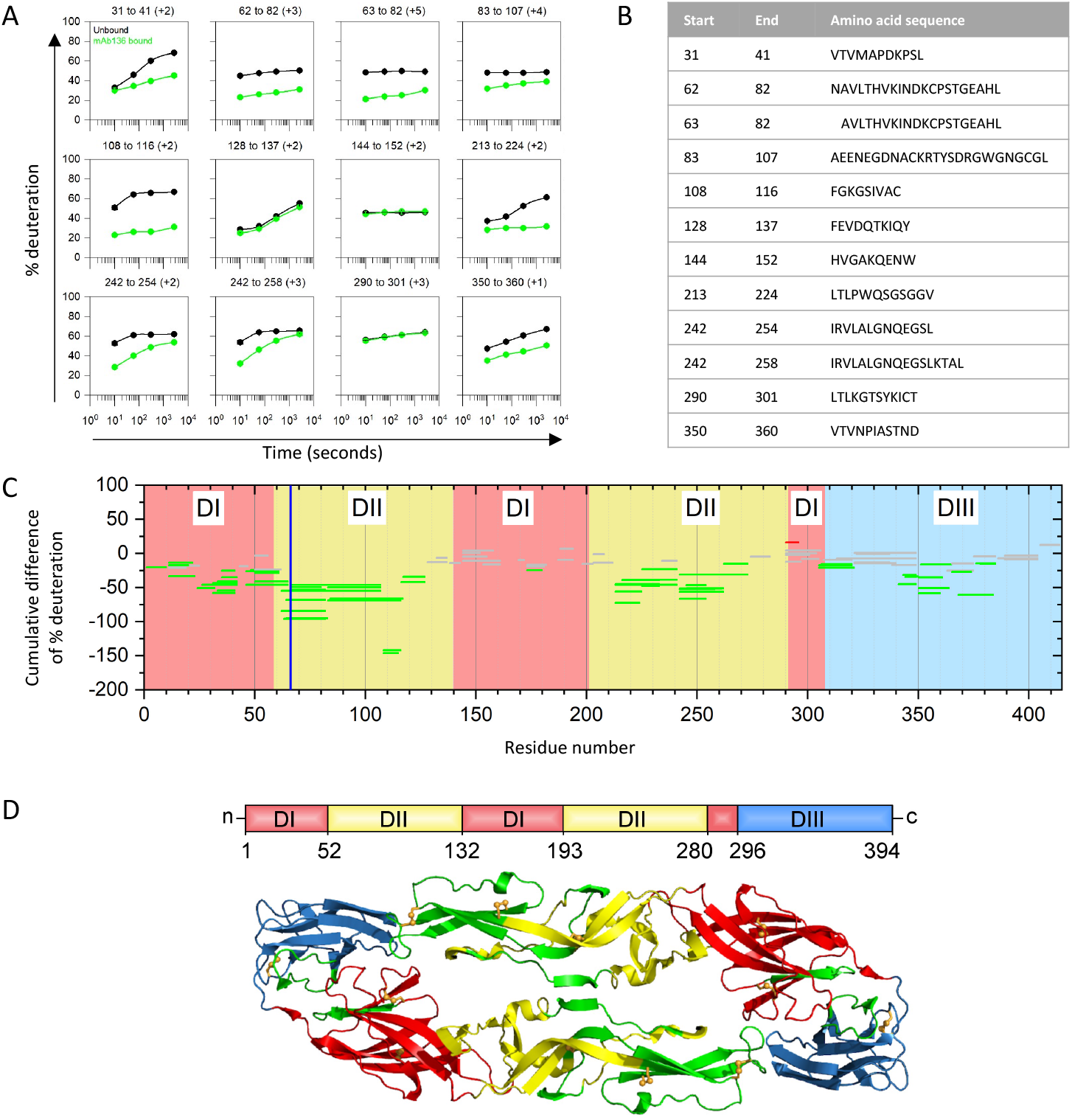
Epitope mapping for Fab YFV-136 by HDX-MS. **A**. Representative kinetic plots for the twelve different peptides showing effects of Fab binding by HDX. Black or green lines are for YFV-E protein in the absence or presence of the mAb136 antibody, respectively. At the top of each panel are the residue numbers and charge states of the peptide. **B**. Sequences and positions of each peptide are given in the table. **C**. Woods plots showing accumulated difference in % deuteration (bound state - unbound state) across all time points for each analyzed peptide. The propagated error for cumulative difference was calculated for each respective peptide, and 99% confidence intervals were calculated. Peptides whose differential exchange exceeds the 99% confidence interval are considered to show significant differences between bound and unbound and involved in binding. Peptides that do not show any significant differences between bound and unbound are in gray whereas protected peptides are in green. The blue vertical line shows the location of H67Y escape mutation identified in studies shown in Figure 4. **D**. The protected peptides are shown in green color on a ribbon representation of the YFV-E dimer. The domains are indicated in red (DI), yellow (DII), or blue (DIII).

### YFV-136 escape mutation studies identify a substitution at H67 that abrogates neutralization capacity

To gain more insight into the epitope of YFV-136 in DII, we selected for neutralization escape variants to identify functionally important interaction residues. To identify mutations in the YFV envelope protein that allow escape from YFV-136 neutralization, we used a real-time cell analysis (RTCA) assay. This high throughput system monitors cell impedance and detects cytopathic effect (CPE) over time, allowing for identification of escape viruses by observation of decreased cell impedance/CPE at late time points after incubating virus with a neutralizing concentration of antibody. For these studies, YFV-17D was incubated with 5 μg/mL of YFV-136 in 16 wells of a 96-well plate. In 13 of 16 wells, complete neutralization and maintenance of cell monolayer integrity was observed throughout the study. However, 3 of 16 wells showed a late CPE phenotype, suggesting selection of variant viruses that subvert YFV-136 neutralization (**Figure 4A**). Supernatants from these wells were harvested and again incubated with 5 μg/mL of YFV-136 on the RTCA instrument to confirm escape. In this second round, CPE developed rapidly, similar to a virus-only control, confirming selection of a population of virus that is refractory to YFV-136 neutralization (data not shown).

**Figure 4.**
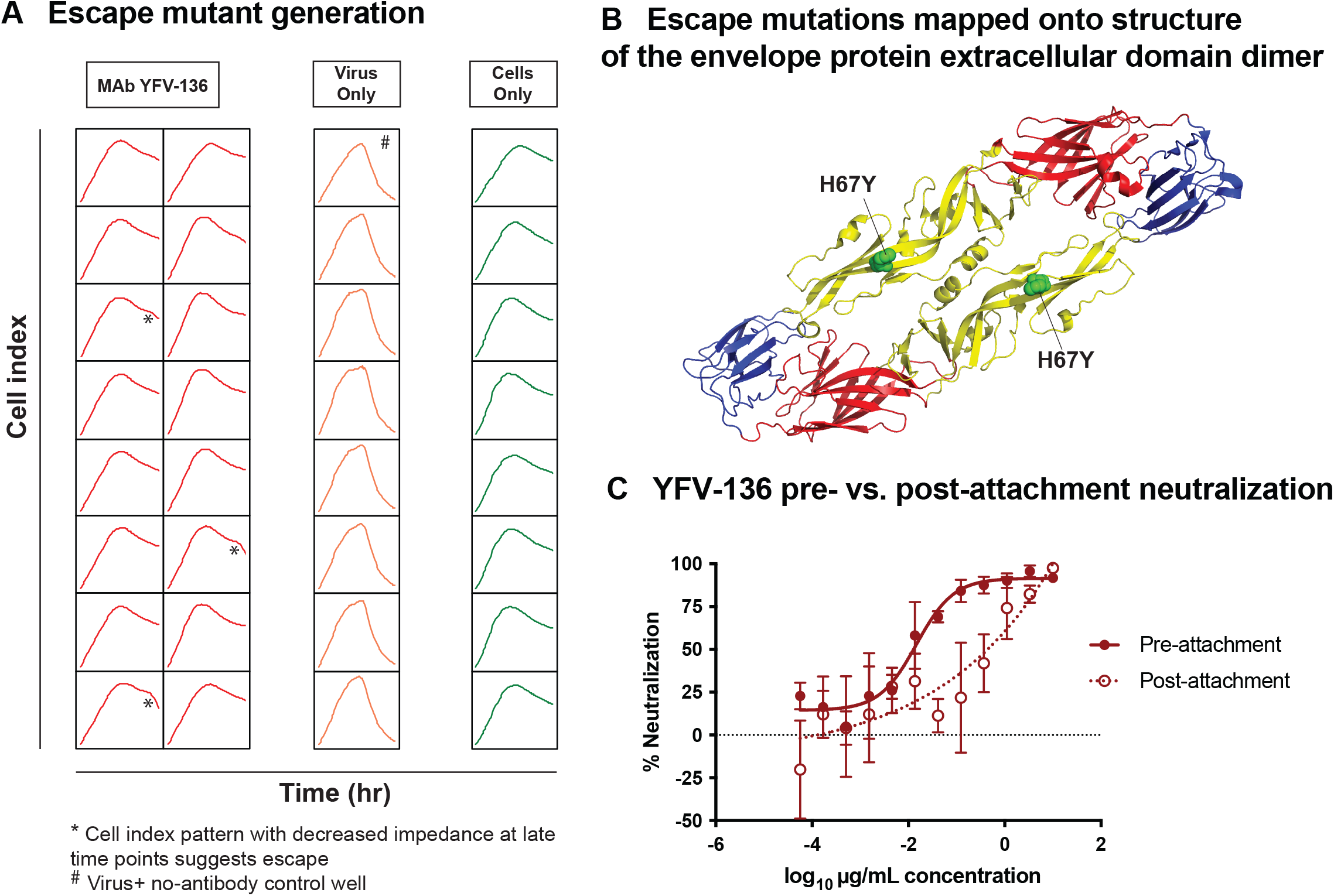
Critical residue for neutralization escape, and mechanism of action studies for YFV-136. **A**. Cell impedance measurements during a first round of YFV-17D escape mutant virus selection. Each box represents the cell impedance within a single well of a 96-well plate as a function of time. The * symbol indicates wells that exhibit a drop in cell impedance at late time points, suggesting viral escape in the presence of 5 μg/mL YFV-136. A single well marked with # was used as a control for cell culture adaptation. * and # wells were propagated once more in culture on the device, and finally in 6-well culture dishes in the presence of 10 μg/mL YFV-136 (or no antibody for #) to allow for viral outgrowth. Viral RNA was isolated and prM and E genes amplified by RT-PCR using primers flanking the prM and E genes. These same primers, and two other primers targeting internal prM and/or E sequences, were used to sequence virus isolated from * and # by Sanger sequencing. These sequences were aligned in Geneious software to identify point mutations. **B**. Escape mutant identified in panel **A** mapped onto the crystal structure of YFV E (PDB 6IW5). Colors denote domain I (red), domain II (yellow), or domain III (blue). **C**. Focus reduction neutralization test of YFV-136 at pre- or post-attachment of virus to host cells. Neutralization values were assessed using a non-linear regression model in Prism software. Two independent experiments were performed in technical triplicate, with data from a single representative experiment shown.

Confirmation of viral escape using RTCA was followed by outgrowth of virus in the presence of 10 μg/mL YFV-136. Viral RNA was isolated, and prM and E genes amplified and sequenced. In all three escape viruses, a single histidine to tyrosine substitution at position 67 on DII of YFV E was identified (**Figure 4B**). This residue in the *b*-strand is conserved across all YFV genotypes, suggesting this escape phenotype likely would be recapitulated in wild-type YFV strains. Overall, escape mutation studies identified a key residue in DII responsible for escape from YFV-136, suggesting this mAb functions by binding an epitope including H67 on DII, consistent with the dominant region of protection from deuteration in the HDX studies.

### MAb YFV-136 neutralizes YFV-17D virus at a post-attachment step

Neutralizing antibodies can target different steps in the viral replication cycle, including, but not limited to, attachment, entry, or egress. To determine the mechanism of action for the most potently neutralizing antibody, YFV-136, we performed pre- and post-attachment neutralization assays **(Figure 4C)**. In the pre-attachment inhibition assay, virus and antibody were pre-mixed prior to addition to Vero cell culture monolayers. In the post-attachment inhibition assay, virus first was incubated at 4°C with cells to allow attachment; afterwards, the excess, unbound virus was washed away, and subsequently antibody was added. YFV-136 neutralized infection in both assays, suggesting that at least part of its inhibitory activity occurs after viral attachment, albeit with some diminished potency.

### MAb YFV-136 protects hamsters from lethal YFV challenge

Because YFV-136 is the most potently neutralizing antibody in our panel, we studied its activity *in vivo*. We first assessed the activity of YFV-136 in a model of YFV disease in Syrian golden hamsters. This model recapitulates many aspects of human YFV infections, including viscerotropism and liver infection and has been used to assess therapeutic efficacy of small-molecules and antibody drugs [24, 30, 31]. Prior to the *in vivo* study, we tested whether the YFV-136 neutralized the hamster-adapted YFV Jimenez strain. MAb YFV-136 exhibited a PRNT_50_ value of 0.5 μg/mL when tested in a PRNT assay in Vero 76 cell monolayers using the hamster-adapted YFV Jimenez strain. Next, animals were administered a 6 × LD_50_ dose of the hamster-adapted Jimenez strain of YFV by an intraperitoneal route. At 3 days post-infection (dpi), 10 animals were treated with 50 mg/kg of YFV-136, and 15 animals with 10 mg/kg of control antibody DENV-2D22 (**Table 1**). Whereas 12 of 15 animals in the control group succumbed to infection, all animals in the YFV-136 group survived the 21-day study (**Figure 5A**). Animals treated with YFV-136 showed transient weight loss after antibody treatment, but quickly recovered and gained weight throughout the remaining course of the study (**Figure 5B**). Viremia was assessed in all animals at day 6 after inoculation. While control mAb (DENV-2D22)-treated animals showed substantial viremia at day 6, animals treated with YFV-136 showed a significant reduction (**Figure 5C**). Finally, we assessed the ability of YFV-136 to prevent YFV-induced liver damage in hamsters by measuring serum alanine aminotransferase (ALT). Whereas animals treated with an isotype control showed markedly increased serum ALT, animals treated with YFV-136 had lower ALT levels **(Figure 5D**), suggesting YFV-136 protects hamsters from hepatic damage induced by the hamster-adapted YFV stain Jimenez.

**Table 1.** The effect of delayed treatment (3 dpi) with mAb YFV-136 in a Syrian golden hamster model of YFV Jimenez strain infection and disease.

| MAb treatment | Dose of mAb given on 3 dpi (mg/kg) | Virus given i.p. on 0 dpi | Alive/total at 21 dpi | Mean day of death ± SD | Mean body weight change<sup>b</sup> (grams ± SD) | Serum virus titer on 4 dpi (CCID<sub>50</sub>/mL ± SD) | Serum alanine aminotransferase test on 6 dpi (mean IU/L ± SD) |
| --- | --- | --- | --- | --- | --- | --- | --- |
| YFV-136 | 50 | YFV<sup>a</sup> | 10/10 | $>21 \pm 0.0^{**}$ | $-5.2 \pm 6.7$ | $4.1 \pm 2.2$ | $114 \pm 26^*$ |
| Isotype control (DENV 2D22) | 10 | YFV<sup>a</sup> | 3/15 | $7.5 \pm 1.4$ | $-8.9 \pm 9.9$ | $5.4 \pm 2.3$ | $214 \pm 120$ |
| Normal controls | -- | Sham | 5/5 | $>21.0 \pm 0.0^{**}$ | $3.0 \pm 2.1^{**}$ | $1.7 \pm 0.0^{**}$ | $80 \pm 3^*$ |

<sup>a</sup> Hamster-adapted YFV Jimenez strain [24].785 <sup>b</sup> Difference between weight on 4 and 5 dpi, representing maximal weight change in this study.786 <sup>\*\*</sup>P<0.001, <sup>\*</sup>P<0.01, as compared to control treatment.

**Figure 5.**
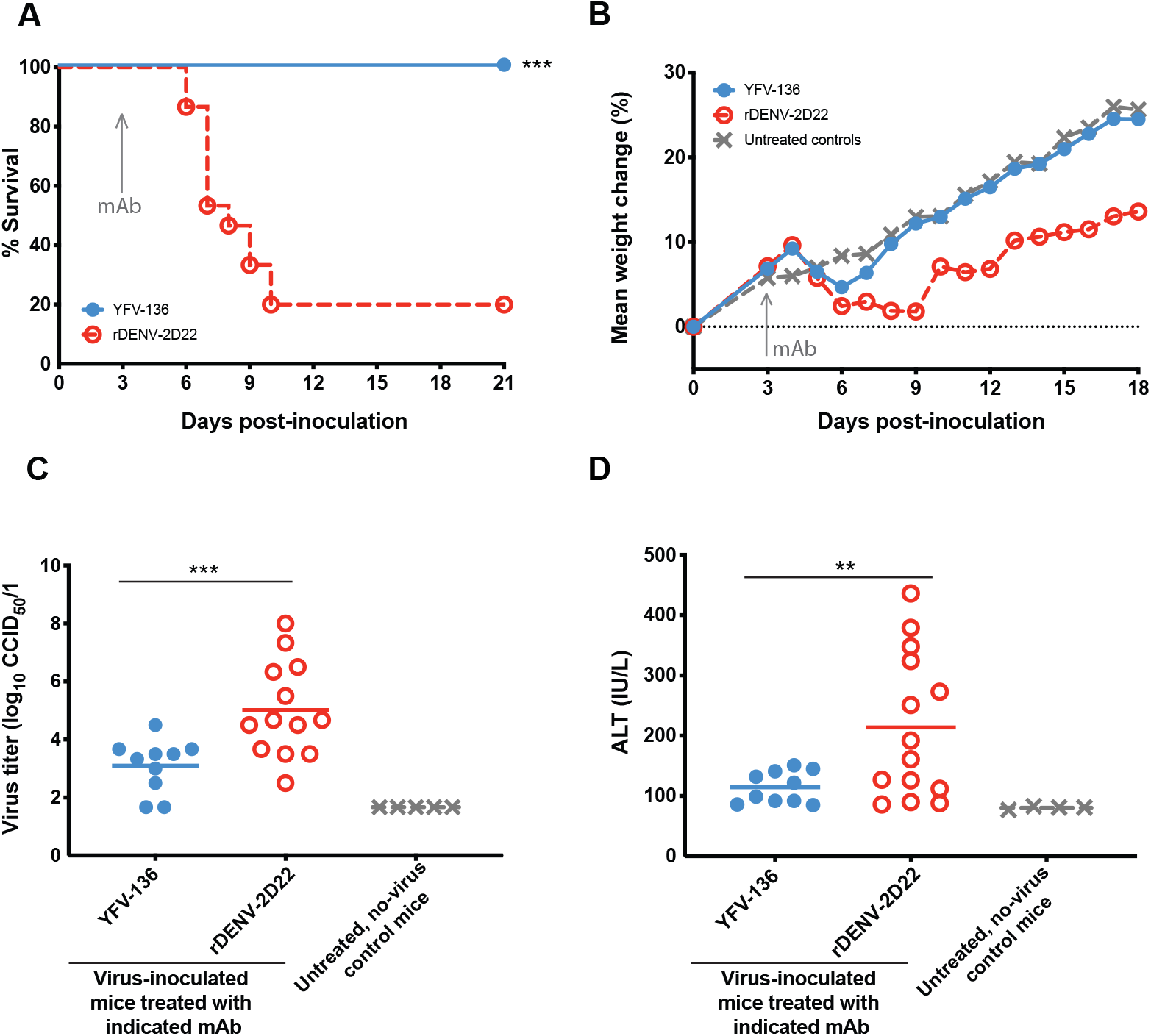
Syrian golden hamster challenge studies to assess YFV-136 therapeutic efficacy. **A**. Kaplan-Meier survival curves of animals (YFV-136, n=10; DENV 2D22 control, n=15; uninfected controls, n=5) treated with 50 mg/kg YFV-136 or 20 mg/kg isotype control 3 days post-inoculation with 200 × CCID_50_ hamster-adapted YFV Jimenez strain. Statistical analysis was performed using a Wilcoxon Log-Rank test. **B**. Weights of YFV-infected animals treated with YFV-136 or isotype control mAb, or uninoculated animals throughout the course of the study. **C**. Serum virus titers (50% cell culture infectious doses [CCID_50_]) were assessed 6 days after virus inoculation. A one-way ANOVA with Dunnett’s multiple comparisons was used to assess statistical significance. **D**. Serum alanine aminotransferase (ALT) levels at 6 days after inoculation was assessed as a proxy for liver damage. A one-way ANOVA with Dunnett’s multiple comparisons post-test was used to assess statistical significance.

### YFV-136 protects humanized mice from lethal YFV challenge

We recently developed a YFV infection model in mice engrafted with human hepatocytes (hFRG mice) [32]. Immunodeficient hFRG mice have three genetic lesions (fumarylacetoacetate hydrolase *[<u>F</u>ah]^-/-^, <u>R</u>ag2*^-/-^, and *Il2r<u>ɣ</u>*^-/-^ on a C57BL/6J background), which together with specifically-timed dietary modifications, facilitate the durable replacement of murine hepatocytes with transplanted human hepatocytes [33]. YFV-infected hFRG mice developed disease that recapitulates many features of YF in humans including massive hepatic infection and injury [32]. Here we tested the therapeutic activity of YFV-136 in this highly susceptible hepatotropic model. hFRG mice were administered a single 10 mg/kg dose of YFV-136 or isotype control mAb (DENV-2D22) at 8 hr after inoculation with 2 × 10^5^ focus forming units (FFU) with *wt* YFV-Dakar (DakH1279), a highly pathogenic West African strain. By 4 dpi, all isotype mAb-treated hFRG mice displayed substantial signs of disease: two were dead, the other three were moribund, and all had lost 15 to 25% of their initial body weight (**Figure 6A-B**). In contrast, hFRG mice that were inoculated with YFV and treated with YFV-136 mAb appeared healthy and exhibited minimal weight loss. The reduced disease observed in the YFV-136-treated group corresponded with significant (~1,000-fold) reductions in YFV burden in the serum and liver (**Figure 6C-D**), and normal levels of liver synthetic function (as measured by the prothrombin time [PT]), hepatocyte damage (alanine aminotransferase [ALT]), biliary function (bilirubin), and detoxification capacity (ammonium) (**Figures 6E-H**). Thus, YFV-136 therapy given 8 hr post-infection was highly protective in the susceptible hFRG mouse model of YFV infection and liver disease.

**Figure 6.**
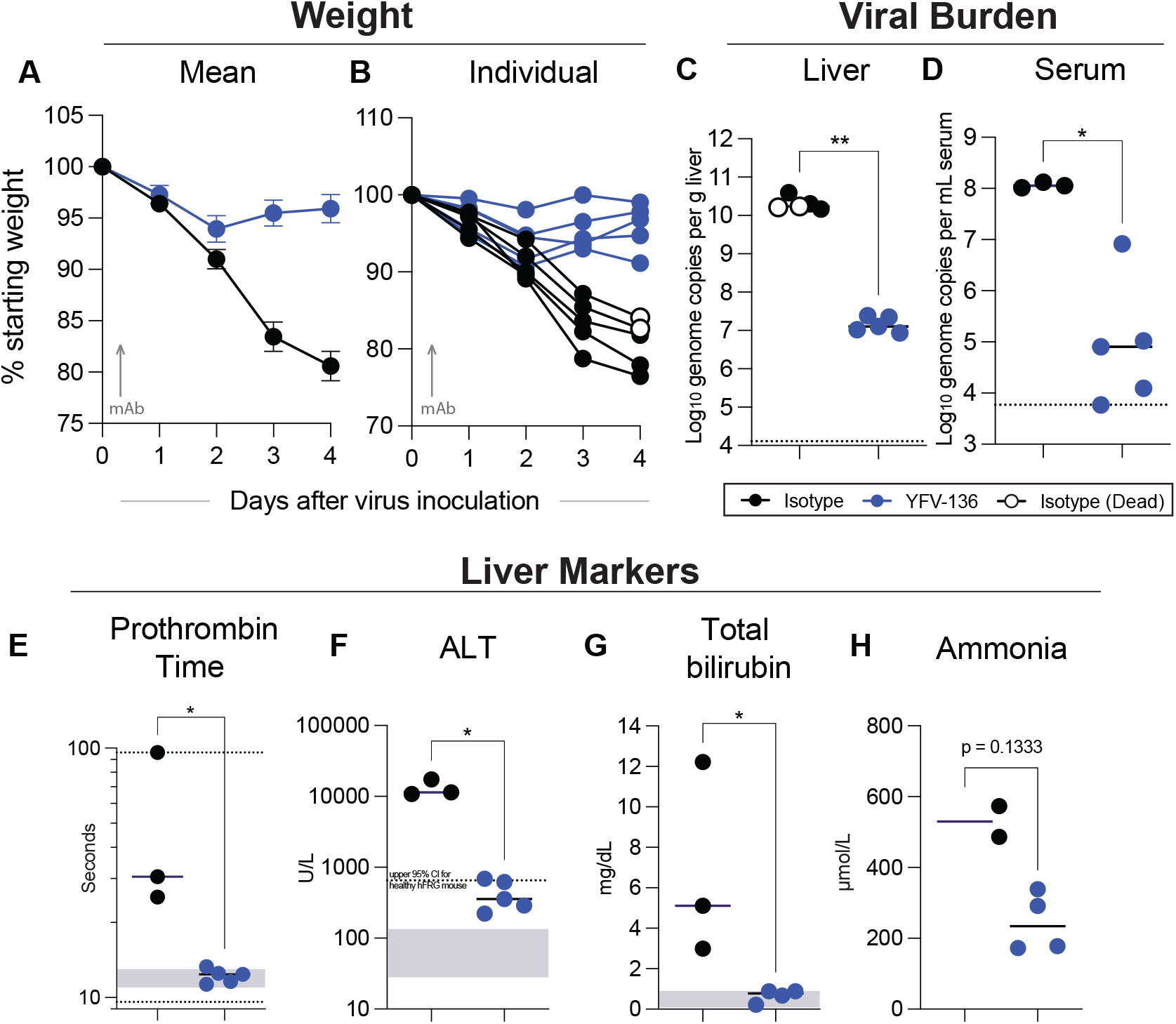
hFRG challenge studies to assess YFV-136 therapeutic efficacy. FRG mice engrafted with human hepatocytes (hFRG) were inoculated with 2×10^5^ focus forming units (FFU) of *wt* YFV-DakH1279. Eight hr later, mice were administered a single 10 mg/kg dose of YFV-136 (n = 5, blue points) or isotype control (n = 5, black points). Note, two of the control mAb-treated YFV-infected mice succumbed to infection at 4 dpi (denoted by open circles), and thus some serum-based measurements were not available. **A-B**. Weight loss showing mean values (**A**) and individual animal profiles (**B**). **C-D**. Viral burden measures at 4 dpi in the liver (**C**) and serum (**D**) as measured by RT-qPCR. **E-H**. Serum biomarkers of hepatic injury at 4 dpi. Samples were tested for prothrombin time (**E**), alanine aminotransferase (ALT, **F**), total bilirubin (**G**), and ammonia (**H**) levels. **C-H**. Mann-Whitney test (*, p<0.05; **, p<0.01). Bars denote median values. Dashed lines in **C and D** indicate the (lower and/or upper) limit of detection of the assay. Grey boxes indicate the reference range for each parameter in mice; for ALT a dashed line is shown to denote the upper limit of “normal” for healthy hFRG mice at baseline.

## Discussion

Yellow fever virus is a re-emerging arbovirus with epidemic potential. While a highly effective live-attenuated vaccine is available for human use, safety and manufacturing concerns warrant new countermeasure development. Here, we isolated a panel of naturally occurring fully human mAbs that bind to the primary target of anti-YFV functional humoral immunity, the E glycoprotein. Two mAbs, YFV-121 and YFV-136, showed neutralization activity against YFV-17D, with YFV-136 showing exceptional potency with IC_50_ < 10 ng/mL. The potency of YFV-136 represents one of the most potent mAbs against YFV ever isolated [23, 26], prompting us to study this mAb in detail. This mAb also neutralizes several wild-type strains of YFV. Both neutralizing mAbs YFV-121 and YFV-136 bind to overlapping antigenic sites as determined by competition binding, suggesting recognition of a shared site of neutralization vulnerability. Antibody escape mutant virus studies identified H67 on DII as a critical residue for the function of YFV-136 within an epitope region in DII identified by HDX-MS studies. We show that YFV-136 functions to inhibit infection at least in part at a post-attachment step in the viral lifecycle. Finally, this mAb was highly efficacious in two different small animal models of YFV infection even when administered after infection, suggesting YFV-136 warrants further development as a therapeutic mAb for use in humans.

Previous studies support the concept that administration of mAbs may be effective in reducing viral load and disease. The murine DII-specific mAb 2C9 and murine-human chimeric mAb 2C9-cIgG exhibited therapeutic activity when administered one day after infection in interferon alpha/beta/gamma receptor-deficient strain AG129 mice challenged with YF 17D-204 vaccine [25] and against virulent YFV infection in an immunocompetent hamster model [24]. A second murine-human chimerized mAb designated 864-cIgG that recognizes DIII and neutralizes the YFV 17D-204 vaccine substrain but did not protect AG129 mice against 17D-204 infection, probably due to its low potency (reported as 10 μg/mL in a 90% plaque reduction neutralization test) [34]. Other investigators have isolated fully human mAbs with similar ultrapotent (<10 ng/mL IC_50_) neutralizing activities [28], although animal protection studies are not reported for these mAbs. One human antibody has been tested in a Phase I clinical trial in which the mAb designated TY014 prevented viremia in 5/5 recipients, whereas only 1/5 placebo recipients lacked viremia at 48 or 72 hr following administration of live yellow fever vaccine (strain YF17D-204; Stamaril)[26]; neutralizing potency and protection in animal studies for this mAb were not reported.

The antigenic site recognized by YFV-136, which lies proximal to the fusion loop (FL) on E protein domain II, has been previously implicated as important for humoral immunity induced by YFV-17D vaccination [28]. This recognition pattern is not restricted to human immune responses, as Ryman *et al*. also showed that some mAbs isolated from mice bind to a site proximal to H67, suggesting broad immunodominance of this site [9]. MAbs characterized in Wec *et al*. display a propensity for pairings of heavy and light chain genes encoded by antibody variable genes *IGHV4-4 + IGLV1-51*, suggesting a public clonotype is elicited by YFV-17D vaccination [28]. Our data complement these findings, as YFV-136 also uses this pairing. The neutralizing activity of YFV-136 is comparable to that of the most potent YFV antibodies reported [28]. It is possible that the efficacy of YFV-17D hinges on its ability to elicit antibodies to this site, since both neutralizing mAbs we isolated from these donors are members of this public clonotype. However, the number of mAbs isolated here is not sufficient to make definitive conclusions in this regard. To date, few studies probing the humoral immune response to YFV have studied the circulating B cells of survivors of natural infection.

The epitope mapping studies using HDX-MS and neutralization escape studies used here suggest that most likely that critical contacts of YFV-136 are focused on the H67 and the region surrounding it on DII. This antigenic site is known in flaviviruses to be a site of vulnerability for potently neutralizing antibodies. For instance, we have observed a similar pattern of binding for the DII-reactive human mAb ZIKV-117 that potently neutralizes the related flavivirus Zika virus [35]; ZIKV-117 also binds detectably to monomeric E protein but it recognizes a quaternary epitope involving two protomers in the virus particle. Higher resolution structural studies are needed to understand the interaction of YFV-136 with virus particles better and to identify a more complete binding footprint for YFV-136.

It is important to note that the antibodies highlighted here bind to monomeric YFV E protein. While we attempted to identify antibodies targeting quaternary structural epitopes only present on virions using a flow-cytometric approach detecting antibodies binding to E protein expressed in YFV-infected cells, these efforts did not identify neutralizing mAbs, and the approach was not explored further. It is likely that functional mAbs targeting sites spanning one or more E protein dimers exist against YFV as has been observed for many flaviviruses. In future work, a multi-pronged approach that employs screens for binding to protein and whole virions, as well as front-end neutralization screens, might help to reveal if immune humans possess some of this rare class of antibodies that exclusively recognize quaternary epitopes on virus particles.

## ACKNOWLEDGEMENTS

We thank Rachel Nargi from Vanderbilt in the Crowe laboratory for technical support with antibody production and purification, and Joseph Reidy and Andrew Trivette for technical support with sequencing. The work of M.P.D was supported by NIH fellowship grant F31 AI152332. J.E.C. is a recipient of the Future Insight Prize from Merck KGaA, which supported this work with a grant. The project described also was supported by R01 AI073755 (M.S.D., D.H.F., and J.E.C.) and the mass spectrometry by R24 GM136766. Vanderbilt University Medical Center has used the non-clinical and pre-clinical services program offered by the Division of Microbiology and Infectious Diseases (DMID) in the National Institute of Allergy and Infectious Diseases (NIAID). The YFV Jimenez strain *in vitro* and hamster studies research were provided under contract number HHSN272201700041I Task Order A11 for the *in vivo* study and 75N93019D00021 Task Order B05 for the *in vitro* studies. The contents of this publication are solely the responsibility of the authors and do not necessarily represent the official views of NIAID or NIH.

## AUTHOR CONTRIBUTIONS

Conceptualization M.P.D., J.R.G., A.L.B., and J.E.C.; Methodology M.P.D., J.R.G., A.L.B, N.K., J.G.J., M.S.D., J.E.C.; Investigation M.P.D., J.R.G., A.L.B., N.K., C.G., J.R., K.M.R., C.A.N., P.N.J., R.E.S., R.G.B., J.G.J.; Resources C.A.N., D.H.F. J.G.J, M.S.D., J.E.C.; Writing – Original Draft M.P.D., J.E.C., Writing – Review & Editing all authors; Supervision M.L.G., J.G.J., D.H.F., M.S.D, J.E.C.; Project Administration M.S.D., J.E.C.; Funding Acquisition M.P.D., M.S.D., J.E.C.

## Methods

### Generation of human mAbs

Blood samples were obtained after written informed consent from four human subjects aged 20 to 47 years who were previously vaccinated with a YFV vaccine prior to travel. The studies were reviewed and approved by the Institutional Review Board of Vanderbilt University Medical Center. Peripheral blood mononuclear cells (PBMCs) were isolated from whole blood and transformed using Epstein-Barr virus (EBV), as previously described [36]. Briefly, transformed B cells were expanded and co-cultured with irradiated human PBMCs in 96-well plates. Cell supernatants were screened by ELISA using recombinant YFV E protein (Meridian Life Sciences). Wells with positive reactivity were fused to a human-mouse myeloma cell line (HMMA 2.5) and plated by limiting dilution in 384-well plates. The resulting hybridomas were cloned by fluorescence-activated cell sorting (FACS) to produce clonal hybridoma cell lines. These clonal hybridoma cells were cultured in T-225 flasks containing serum-free medium, and mAb was purified from spent medium by affinity chromatography on an ÄKTA™ pure Fast Protein Liquid Chromatography (FPLC) instrument (Cytiva).

### Recombinant antibody expression and purification

For animal studies, large-scale recombinant antibody production of YFV-136 was performed. RNA was isolated from YFV-136 hybridoma line, and heavy and light chain genes were amplified using 5’RACE (Rapid Amplification of cDNA Ends) and sequenced using a Pacific Biosciences Sequel instrument. Variable regions of YFV-136 were cloned into a monocistronic full-length, human IgG1 expression vector (Twist Biosciences) for recombinant production. This expression vector then was transfected transiently into ExpiCHO cells for 7 to 8 days. Cell supernatants were harvested and filtered through 0.45-μm filters prior to purification. Purification was performed on an ÄKTA™ pure Fast Protein Liquid Chromatography (FPLC) instrument (Cytiva) using HiTrap MabSelect SuRe columns (Cytiva) as described above for hybridoma-derived mAbs.

### ELISA binding of mAbs to YFV E protein

384-well plates were coated with 2 μg/mL of YFV E protein (Meridian Life Science) at 25 μL/well and incubated overnight at 4°C. Plates then were washed and blocked using Dulbecco’s PBS with Tween 20 (DPBS-T) containing 2% milk and 1% goat serum for 1 hr at room temperature. Following a wash step, serial dilutions of antibody in DPBS were added to plates and incubated for 1 hr at room temperature. To detect bound antibodies, alkaline phosphatase conjugated goat anti-human IgG diluted 1:4,000 in DPBS-T containing 1% milk and 1% goat serum was added to plates for 1 hr at room temperature and developed using AP substrate tablets diluted in 1 M Tris, 0.3 mM magnesium chloride. Plates were developed in the dark for 1 hr and read on a BioTek plate reader at 405 nm. Binding curves were interpolated in Prism software (GraphPad) using a non-linear regression analysis.

### YFV-17D focus reduction neutralization test (FRNT)

A focus reduction neutralization test was performed as previously described with minor amendments. Briefly, 96-well plates were seeded with Vero cells at 2.5 × 10^4^ cells/well and incubated overnight. The following day, serial dilutions of antibody were mixed with 10^2^ FFU YFV-17D and incubated at 37°C for 1 hr. 30 μL/well of virus/antibody mixture then was added to Vero cell culture monolayers and incubated at 37°C for 1 hr. Without washing, 110 μL per well of overlay containing a 1:1 mixture of 2.4% methylcellulose and 2× Dulbecco’s Modified Eagle Medium (DMEM) with 4% FBS was added to plates, which then were incubated for 72 hr at 37°C in 5% CO_2_. To stain foci of virus infection, cells were fixed with 1% paraformaldehyde for 1 hr at room temperature, washed, and permeabilized using permeabilization buffer (0.1% saponin, 0.1% BSA in DPBS) for 10 min. Cells then were stained with 1 μg/mL of pan-flavivirus murine mAb 4G2 in permeabilization buffer for 1 hr at room temperature. After two washes, goat anti-mouse IgG-horseradish peroxidase (Southern Biotech) diluted 1:1,000 in permeabilization buffer was added to cells and incubated for 1 hr at room temperature. Foci were developed using TrueBlue peroxidase and counted using a spot counter instrument (ImmunoSpot; CTL). Foci counts were normalized to that of a virus-only control, and neutralization curves were interpolated in Prism software using a non-linear regression analysis.

### Wild-type and 17D YFV strain FRNT

FRNT was performed as described above, with the following exceptions: 100 μL containing 200 FFU of virus was mixed with 100 μL of serially-diluted mAb and incubated at 37°C for 1 hr. 100 μL of virus:mAb mixture was then added to Vero cells in 96-well plate format and incubated at 37°C for 1 hr, and followed by addition of overlay and incubation at 37°C for two days. Cells were then fixed with 4% paraformaldehyde (final concentration) for 30 min, permeabilized, stained, and analyzed as described above.

### Pre- and post-attachment neutralization of YFV-17D

Pre- and post-attachment neutralization assays were performed as previously described [37]. For pre-attachment studies, 600 FFU YFV-17D was mixed with serial dilutions of antibody for 1 hr at 37°C. Cells and virus/mAb mixtures were then pre-chilled for 15 min prior to addition of mixtures to cell monolayers for 1 hr at 4°C. Cells then were washed three times and incubated with pre-warmed DMEM for 15 min prior to addition of methylcellulose overlay containing DMEM. For post-attachment studies, cell monolayers first were incubated with virus for 1 hr at 4°C. Cells then were washed and incubated with serial dilutions of antibody for 1 hr at 4°C. Excess antibody then was washed off, and cells were incubated for 15 min at 37°C with DMEM prior to addition of overlay. Foci were enumerated as described for the focus reduction neutralization test described above.

### Biolayer interferometry competition-binding assay

Competition-binding studies were performed using a biolayer interferometry instrument (FortéBio Octet® HTX). HIS1K sensortips were pre-incubated in kinetics buffer (Pall) for 10 min. After a 60 sec baseline step, his-tagged YFV E protein (Meridian Life Science) was associated to the tips at 5 μg/mL for 60 sec. Readings again were set to baseline for 60 sec, followed by association of the first antibody at 25 μg/mL for 600 sec to achieve complete saturation. Tip readings again were baselined, and then tips were dipped into wells containing a second antibody at 25 μg/mL for 180 sec. Data were analyzed using FortéBio data analysis software. Data from all steps were normalized to a buffer-only control, and antibodies were grouped using a Pearson correlation statistical analysis.

### Hydrogen-deuterium exchange (HDX) mass spectrometry

#### Soluble recombinant E protein used for HDX studies

A synthetic DNA fragment encoding residues 123 to 680 (VTLV…EGSS) of yellow fever virus strain 17DD-Brazil (GenBank entry AAZ07885.1) E protein was inserted downstream of a modified human IL-2 signal peptide (MARMQLLSCIALSLALVTNSV). The construct was also modified at the C-terminus of the envelope region to contain a small linker, a TEV protease site, and a 6-HIS tag (GSTGGSENLYFQGHHHHHH). The fusion construct was inserted into an AgeI-NotI-cut pFM1.2R vector [38] by Gibson assembly to lie downstream of the CMV promotor. Recombinant E protein was produced by transient transfection of Expi293F cells using a ExpiFectamine 293 Transfection Kit (Thermo Fisher Scientific). Cell supernatants were harvested 4 days after transfection and then concentrated before exchange into 2× PBS at pH 6.5 and finally into 2× PBS at pH 8.0. The soluble recombinant E protein was recovered by 6-HIS affinity chromatography on Ni-NTA agarose (G-Biosciences) and purified by size exclusion chromatography on a Superdex S200 Increase column (Cytiva).

#### Peptide mapping

To prepare for acquisition and analysis of HDX data, the YFV-E protein was digested with two acid proteases (immobilized pepsin followed by immobilized acid protease from fungal-XIII) to achieve better sequence coverage. To effectively reduce the protein and denature it, a quench solution tris (2-carboxyethyl) phosphine hydrochloride, TCEP) and guanidine hydrochloride, GdnHCl was added. The quenching conditions were 1:1 dilution of HDX reaction volume (100 μL) with quench buffer containing 500 mM TCEP and 4 M GdnHCl, pH 2.4 (resulting in a 250 mM TCEP, 2 M GdnHCl and pH 2.6), 3 min incubation, 25°C. A peptide map (in triplicate) of the digest was generated by LC-MS/MS using a Maxis-II-HM mass spectrometer (Bruker Daltonics, Billerica, MA). YFV-E protein (100 pmol) was injected in the LC-MS system where the protein was digested, and the resulting peptides were captured and desalted by C-8 trap column (2.1 × 20 mm, Zorbax Eclipse XDB-C8 trap (Agilent) followed by loading on to C-18 column (2.1 × 50 mm in size, 2.5 μm Xselect-CSH from Waters, Milford, MA) and elution into the mass spectrometer. The mass spectrometer was operated in a data-dependent fragmentation mode with monitoring the high abundance peptides. Data were analyzed by Byonic™ (Protein Metrics, Santa Carlos, CA, USA) for sequencing and accurate precursor mass (± 5 ppm), and the peptides also were curated manually.

#### HDX experiment

YFV-E was equilibrated without or with antibody (1:2 antibody) in phosphate buffered saline in H_2_O (PBS, pH 7.4) overnight at 4°C and reequilibrated at 25°C for 30 min before starting the HDX experiment. The exchanged-in with D_2_O (PBS prepared in D_2_O) in the absence (10 μM YFV-E) or presence of antibody (20 μM antibody, (1:2 antigen: antibody ratio) was initiated by diluting the protein solutions (10 μL) by 10-fold with D_2_O in PBS at 25°C (90 μL, pH 7.4). HDX was measured at 0 (undeuterated control), 10, 60, 300, and 2,700 s at 25 °C. For the undeuterated control, the conditions were the same except the added buffer solution was H2O instead of D_2_O. The HDX was quenched by adding an equal volume (100 μL) of quench buffer equilibrated at 25°C followed by mixing and incubation at 25°C for 3 mins. The quenched sample was digested by passing it through a custom-packed column (2 mm x 20 mm) of immobilized pepsin beads followed by a column of immobilized Fungal XIII beads (2 mm x 20 mm) at 200 μL/min flow rate. The resulting peptides thus generated were captured and desalted on a C-8 column by washing with 0.1% formic acid in water for 4.7 min. Desalted peptides were loaded on a C-18 analytical column where peptides were separated using a gradient of acetonitrile (ACN) in 0.1% formic acid (most peptides eluted during the linear part of gradient from 5 min (4% ACN) to 15 min (40% ACN)). To minimize back exchange, the trap and analytical columns were kept in an ice slush while protease columns were at room temperature. The isotope distributions of the exchanged peptides were measured with a Maxis-II-HM mass spectrometer (MS only mode) for duplicate samples.

#### HDX data analysis

LC-MS HDX data acquisition (retention time, isotopic distribution and observed *m/z*) was directed by the peptide map, and data were analyzed by HDExaminer (v 2.5.0, 64-bit, Sierra Analytics). The maximum deuterium level was set to 90%, and the data displayed as kinetic plots for each peptide for HDX. Only those peptides that provided good signal-to-noise ratio at all the time points and for both the states were included in the analysis. Ultimately, 107 unique peptides covering 95% sequence of YFV-E protein were analyzed. Average peptide length was 14 amino acids and average residue level redundancy was 4. To elucidate those regions where HDX changed with statistical significance upon antibody binding, a mean cumulative difference (bound - unbound) across all the time points for each peptide was calculated and plotted as a Woods plot. To identify significant differences upon binding, the propagated error for cumulative percent deuteration difference for each peptide was calculated using the standard error of mean, and a 99% confidence interval was determined (2 degrees of freedom, two tail distribution). Peptides that showed no change are marked in gray and peptides that showed significant differences between bound and unbound were highlighted based on protection (green) or exposure (red).

### Generation and analysis of YFV-17D escape mutations

In a U-bottom 96 well plate, 25 μL mAb YFV-136 IgG protein at 5 μg/mL was pre-mixed with 25 μL YFV-17D virus [39] diluted 1:10 (~5,000 FFU) in DMEM without FBS and incubated for 1 hr at 37°C. This procedure was done in 16 separate wells within the 96-well plate. Virus also was mixed with DMEM alone and passaged throughout the study to control for substitutions that arise from cell culture adaptation. 50 μL virus/antibody mixture and controls were added to HuH-7.5 cell culture monolayers in 96-well ePlates (Agilent) and incubated for 1 hr at 37°C. 100 μL of DMEM containing 5% FBS was then added to each well, plates placed back on an xCELLigence instrument (Agilent), and cell monolayers were monitored for delayed CPE. Cell supernatants in wells with a delayed CPE phenotype, as well as a virus-only control, were subjected to a repeat of this assay to confirm viral escape. Once escape was confirmed, 6-well plates containing confluent Huh7.5 cell monolayers were inoculated with 100 μL/well escape virus or a virus control in the presence of 10 μg/mL of YFV-136 for outgrowth of escape virus. Virus was harvested from 6-well plates, and RNA isolated using the Qiagen virus RNA isolation kit. E and prM genes from isolated RNA were reverse-transcribed to cDNA and PCR-amplified using primers flanking the prM and E genes (One-step RT-PCR kit). Genes then were sequenced by GENEWIZ using overlapping primers that give coverage across prM and E. The control virus sequence was aligned to the 17D reference genome sequence to confirm that mutations did not result from adaptation to cell culture.

### Syrian golden hamster challenge studies

The Syrian golden hamster model used for these studies has been described previously [24]. 30 female Syrian golden hamsters (LVG/Lak strain) supplied by Charles River were used. Hamsters were randomized by weight to experimental groups and individually marked with ear tags. For challenge studies, hamsters were challenged at day 0 with 200 of 50% cell culture infectious doses (CCID_50_) hamster-adapted YFV Jimenez strain by bilateral intraperitoneal injections in a total of 0.2 mL. At 3 days post-virus inoculation, hamsters were dosed at 50 mg/kg of recombinant YFV-136 (rYFV-136) (1 mL total volume) or 10 mg/kg of rDENV-2D22 control and monitored for weight loss and clinical manifestations for 21 days. Blood samples were taken at days 4 and 6 to assess viremia and ALT. Any surviving animals were humanely euthanized at the experimental endpoint.

### Measurement of hamster serum aminotransferase

ALT (SGPT) reagent (Teco Diagnostics, Anaheim, CA) was used, and the protocol was altered for use in 96-well plates. Briefly, 50 μL aminotransferase substrate was placed in each well of a 96-well plate, and 15 μL of sample was added at timed intervals. The samples were incubated at 37°C, after which 50 μL color reagent was added to each sample and incubated for 10 min as above. A volume of 200 μL of color developer was next added to each well and incubated for 5 min. The plate was then read on a spectrophotometer, and aminotransferase concentrations were determined per manufacturer’s instructions.

### 50% cell culture infectious doses (CCID_50_) assays to assess hamster viral burdens

Virus titer was quantified using an infectious cell culture assay in which a volume of either tissue homogenate or serum was added to the first tube of a series of dilution tubes. Serial dilutions were made and added to Vero cell monolayer cultures. Ten days later, CPE was used to identify the endpoint of infection. Four replicates were used to calculate the CCID_50_ of virus per mL of plasma or gram of tissues.

### Mouse studies

All mouse experiments were conducted under a Washington University School of Medicine Institutional Animal Care and Use Committee approved protocol in compliance with the Animal Welfare Act. Female hFRG mice were generated by Yecuris Corporation and maintained according to their specific care and use guidelines (see [32] for additional details). Only mice with human albumin levels of ≥ 5.0 mg/mL in plasma (indicative of ≥70% engraftment) were used for this study. hFRG mice were continued on their regular diet of PicoLab High Energy Mouse Diet, 5LJ5 chow (LabDiet) and 3.25% w/v dextrose-water during infection experiments. Mice were inoculated with via retro-orbital injection of 50 μL containing 2 ×10^5^ FFU *wt* YFV-DakH1279, obtained from the World Reference Center for Emerging Viruses and Arboviruses and passaged once in Vero-CCL81 cells. Antibodies were given as a single treatment of 10 mg/kg dose, diluted in PBS in a 100 μL total volume and given by the i.p. route, 8 hrs post-infection. The DENV2D22 mAb was used as a control. Euthanasia was performed via ketamine overdose and thoracotomy. Blood was collected via aspiration from the cardiac chambers and PT was measured on a Coagucheck (Roche). Perfusion of the entire vascular tree was then performed with saline prior to liver collection. After centrifugation, serum was mixed 1:9 with 10% Triton X-100 in HBSS (1% final volume of Triton-X-100), then incubated at room temperature for 1 h to inactivate infectious virus [32]. ALT, bilirubin, and ammonia were analyzed using the Catalyst Dx Chemistry Analyzer (IDEXX Laboratories). Some specimens required dilutions to achieve an ALT value within the analytical measurement range; in these instances, dilution was performed in HBSS and the value was corrected by the final dilution factor. RNA extraction and viral load analysis was performed as described previously [32] using the KingFisher Flex (Thermo Fisher) and the TaqMan RNA-to-CT 1-Step Kit (Thermo Fisher) on the QuantStudio 6 Flex Real-Time PCR System with the following primers: F: 5’-AGGTGCATTGGTCTGCAAAT-3’; R: 5’-TCTCTGCTAATCGCTCAAIG-3’; P: 5’-/56-FAM/GTTGCTAGGCAATAAACACATTTGGA/3BHQ_1/-3’.

